# A Methodology for Evaluating the Performance of Alerting and Detection Algorithms Running on Continuous Patient Data

**DOI:** 10.1101/182154

**Authors:** Larry J. Eshelman, Minnan Xu-Wilson, Abigail A. Flower, Brian Gross, Larry Nielsen, Mohammed Saeed, Joseph J. Frassica

## Abstract

**Objectives:** Clinicians in the intensive care unit (ICU) are presented with a large number of physiological data consisting of periodic and frequently sampled measurements, such as heart rate and blood pressure, as well as aperiodic measurements, such as noninvasive blood pressure and laboratory studies. Because this data can be overwhelming, there is considerable interest in designing algorithms that help integrate and interpret this data and assist ICU clinicians in detecting or predicting in advance patients who may be deteriorating. In order to decide whether to deploy such algorithms in a clinical trial, it is important to evaluate these algorithms using retrospective data. However, the fact that these algorithms will be running continuously, i.e., repeatedly sampling incoming patient data, presents some novel challenges for algorithm evaluation. Commonly used measures of performance such as sensitivity and positive predictive value (PPV) are easily applied to static “snapshots” of patient data, but can be very misleading when applied to indicators or alerting algorithms that are running on continuous data. Our objective is to create a method for evaluating algorithm performance on retrospective data with the algorithm running continuously throughout the patient’s stay as it would in a real ICU.

**Methods:** We introduce our evaluation methodology in the context of evaluating an algorithm, a Hemodynamic Instability Indicator (HII), for assisting bedside ICU clinicians with the early detection of hemodynamic instability before the onset of acute hypotension. Each patient’s ICU stay is divided into segments that are labelled as hemodynamically stable or unstable based on clinician interventions typically aimed at treating hemodynamic instability. These segments can be of varying length with varying degrees of exposure to potential alerts, whether true positive or false positive. Furthermore, to simulate how clinicians might interact with the alerting algorithm, we use a dynamic alert supervision mechanism which suppresses subsequent alerts unless the indicator has significantly deteriorated since the prior alert. Under these conditions determining what counts as a positive or negative instance, and calculations of sensitivity, specificity, and positive predictive value can be problematic. We introduce a methodology for consistently counting positive and negative instances. The methodology distinguishes between counts based on alerting events and counts based on sub-segments, and show how they can be applied in calculating measures of performance such as sensitivity, specificity, positive predictive value.

**Results:** The introduced methodology is applied to retrospective evaluation of two algorithms, HII and an alerting algorithm based on systolic blood pressure. We use a database, consisting of data from 41,707 patients from 25 US hospitals, to evaluate the algorithms. Both algorithms are evaluated running continuously throughout each patient’s stay as they would in a real ICU setting. We show how the introduced performance measures differ for different algorithms and for different assumptions.

**Discussion:** The standard measures of diagnostic tests in terms of true positives, false positives, etc. are based on certain assumptions which may not apply when used in the context of measuring the performance on an algorithm running continuously, and thus repeatedly sampling from the same patient. When such measures are being reported it is important that the underlying assumptions be made explicit; otherwise, the results can be very misleading.

**Conclusion:** We introduce a methodology for evaluating how an alerting algorithm or indicator will perform running continuously throughout every patient’s ICU stay, not just for a subset of patients for selected episodes.

## INTRODUCTION

Predictive algorithms based on streaming patient vital signs are gaining popularity as more sensors become available for monitoring in various care settings, i.e. in the intensive care unit (ICU), general ward, and at home (Figure 1). These algorithms aim to integrate large volumes of data into actionable information for care providers. We have been developing and evaluating such algorithms for the intensive care environment to help clinicians detect cardiovascular deterioration in its early stages, enabling them to direct attention to those patients who may benefit from it most. [1] [2] These algorithms use data that is frequently collected, such as heart rate and blood pressure, and the algorithms are often running continuously throughout a patient’s ICU stay. There are many ways to evaluate such algorithms. One common method is to take a static, “snapshot” approach, looking at discrete data points chosen from stable and unstable (i.e., deteriorating) segments. For example, one might sample stable segments from a randomly chosen time points and sample unstable segments from a point shortly (e.g., one hour) before the point of deterioration. Typically, one will examine the feature value one is interested in for a time window at these time point, e.g., a three-hours window prior to the time point, and choose the most recent value or the median value, or some other representative value. These representative values and then treated very much like one would treat the results of a diagnostic tests (such as a PSA test for prostate cancer). Static analysis techniques such as receiver operator curve (ROC) analysis, and performance measures such as sensitivity, specificity, positive predictive value (PPV) and negative predictive value (NPV) can then be applied. However, it is important to recognize how detection algorithms running in a patient monitoring environment differ from diagnostic tests, In the case of diagnostic tests, usually a single test is performed per subject, determining whether the subject is in the stable/healthy group (e.g., doesn’t have prostate cancer) or the unstable/unhealthy group (e.g., has prostate cancer). In a patient monitoring setting, on the other hand, these tests are, in effect, being performed repeatedly, sometimes as often as new data is coming in, which may be continuously. The amount of data available for analysis vary drastically from patient to patient: some patients have been in the ICU for days where others may have only just arrived. In this setting a continuously running algorithm based on high frequency data is aimed at detecting unstable “states” rather than unstable individuals. In fact, the same patient may go from a stable state to an unstable one to another stable one, and so on with some frequency. These state transitions are not represented by a single test with a cut off for positive and negative result but a number of physiological “tests” running simultaneously which represent the patient’s changing physiologic state. During a state transition a patient might be in a stable condition according to the test, an unstable state or a transitional state between the two extremes.

**Figure 1:**
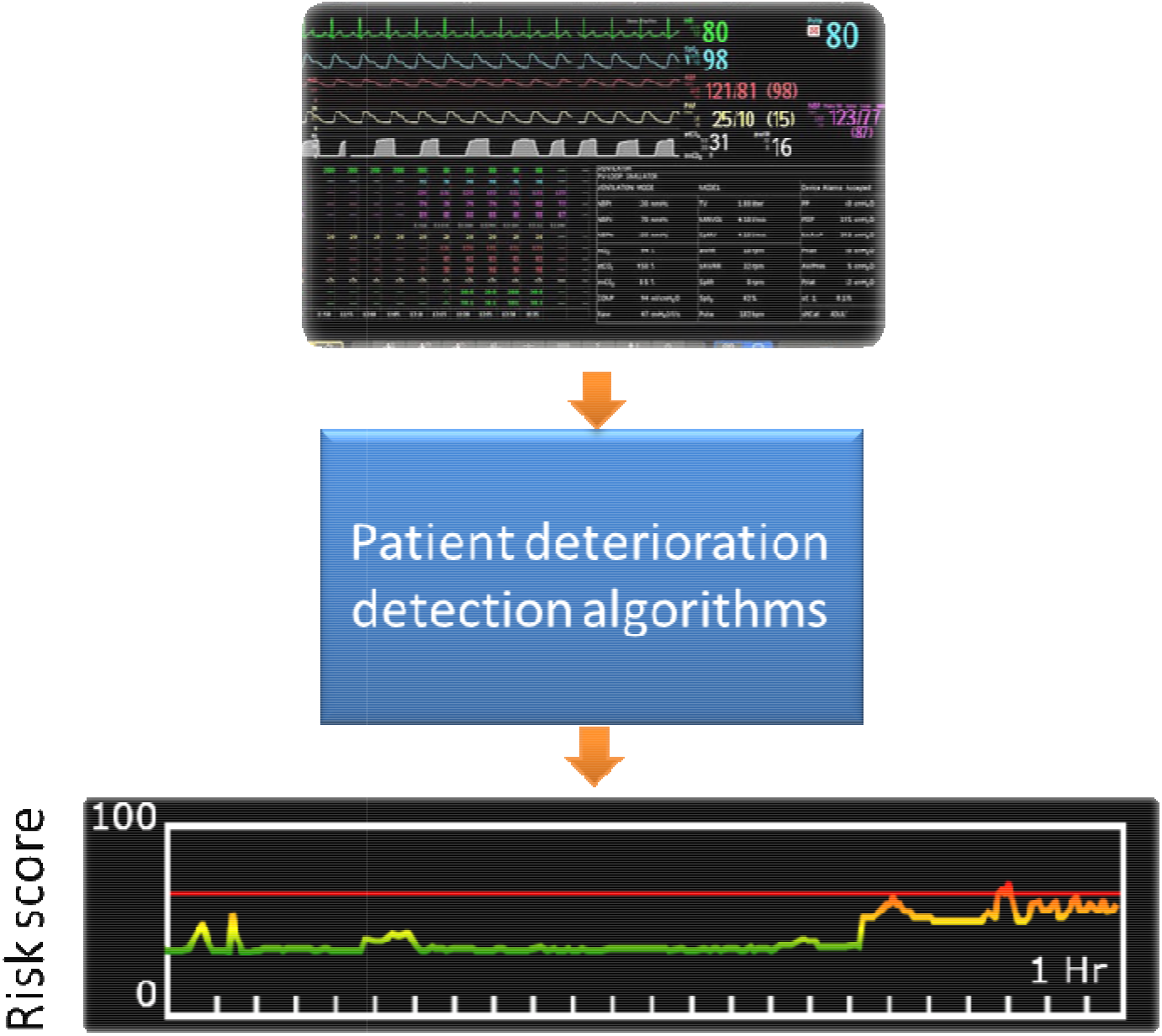
Patient deterioration algorithms are becoming more common as more sensors are available both in the hospital and at home. These algorithms typically output a risk score or sound an alert. Quantifying the performance of these algorithms is challenging with traditional methods developed for diagnostic tests in which a single test is applied once per subject. New methods are needed to characterize how continuously monitoring algorithms perform in the face of real world data where patient health status is often uncertain, data recording can be of varying lengths, and prevalence of the events that the algorithms are trying to detect is brief in time. Furthermore, evaluation methods should capture how clinicians might interact with the algorithm.

Given these differences, it is challenging for traditional tests of performance (as applied to diagnostic tests) to accurately capture how an algorithm based on high frequency data would do in the real world. One challenge is how to apply consistent definitions of what are positives and negatives. Sensitivity, specificity, PPV and NPV are defined in terms of the number of true positives (TP), number of true negatives (TN), number of false positives (FP), and number of false negatives (FN) (see Table 1). Yet these terms are meaningful only to the extent that what we are counting is clearly defined and a consistent set of definitions is used for all terms, e.g., a TP is counted the same way when used for determining sensitivity as when determining the PPV. However, in the context of high frequency data-based alerts inconsistency is often introduced. For example, when calculating PPV it is natural to count alerts, as PPV is typically thought of as the number of true alerts divided by the total number of alerts. But when calculating sensitivity, it is natural to count states, where sensitivity is the number of unstable states correctly identified by at least one alert divided by total number of unstable states. A TP has different meanings in these two calculations. In one case alerts are being counted, in the other states.

**Table 1:**
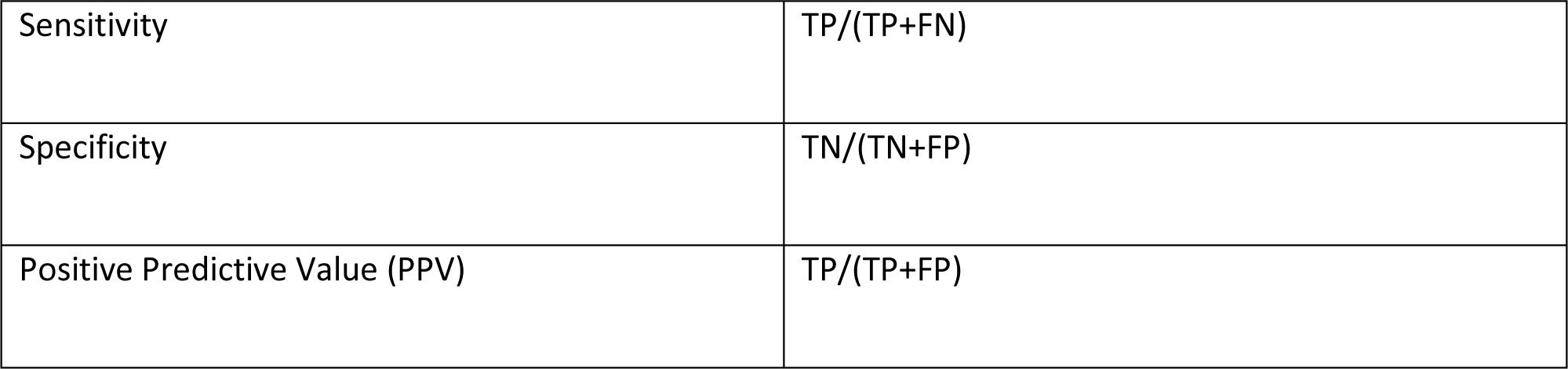
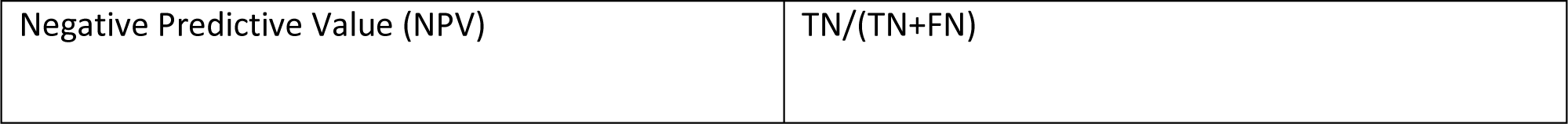
Definitions of sensitivity, specificity, PPV, and NPV in terms of the number of true positives (TP), true negatives (TN), false positives (FP), and false negatives (FN).

Specificity and NPV are even less straightforward since their calculations make use of the number of negative instances without alerts. A negative instance is simple when taking a snapshot approach such as diagnostic tests, but it is not obvious how one should count the negatives when the algorithm is running continuously utilizing high frequency data. Does a stable episode without any alerts count as a single true negative? Or does a stable episode without any alerts that is 3 days long get more true negative credit than one that is only half a day long? These are the sort of questions that need to be asked when analyzing the performance of a detection or predictive algorithm that is repeatedly sampling from continuous data streams. To simply analyze performance as one would a diagnostic test that is performed infrequently is likely to oversimplify the problem and lead to a misleading analysis. In particular, there are four sources of potential confusion:

(1) Snapshots are often not representative samples of patients nor their state of health, e.g., they may only sample clear-cut episodes, and so will not reflect the actual performance for this type of test in realistic conditions.
(2) Often the definitions of TP, FP, TN, and FN are inconsistent with each other, i.e., some based on counting alerts and others patient states.
(3) Snapshots of stable and unstable instances often do not reflect the true prevalence of the unstable instances. Prevalence refers to the occurrence of unstable states in either duration or frequency. Unless prevalence is taken into account, the result PPV and NPV calculations, are likely to be wrong.
(4) Snapshots may not reflect alerting dynamics. Dynamics refer to how alerts maybe interpreted by clinicians and how alerts may go above and below threshold crossings many times.

In the next section we will elaborate on these issues and describe how they can be addressed.

## METHODS

We will explain our method for evaluating a continuously running detection or predictive algorithm using as an example an algorithm that was developed for the prediction of hemodynamic instability. Hemodynamic instability, which can result in acute hypotension, affects approximately 10 to 30 percent of patients in the ICU, can have devastating consequences, and often is detected well after blood pressure (BP) has reached critically low levels (e.g., systolic BP < 90 mmHg or mean arterial pressure < 65 mmHg). The early detection of hemodynamic instability episodes and timely initiation of appropriate corrective intervention can significantly improve patient outcome [3, 4, 5]. We created an algorithm, hemodynamic instability indicator (HII), that can assist bedside clinicians with the early prediction of hemodynamic instability in the ICU before the onset of acute hypotension. HII combines vitals measurements (heart rate and blood pressure) into a vitals index and laboratory measurements (white blood cell count, blood urea nitrogen, hematocrit, albumin, and bicarbonate), which are measured less frequently, into a labs index. It then integrates the two indices into a composite index with a range from 1 to 100 (a higher score indicating a higher risk of instability). [6]

For the development and evaluation of HII we used patient data made available to us by the eResearch Institute [7]. Here we will focus on the evaluation of the algorithm when running continuously. Although the data included records from 105,000 patients’ cared for at 50 hospitals, we refined our dataset to include only those hospitals that reported fluids administered at least hourly so that we could better gauge each patient’s therapy. From these hospitals, we used only data from patients who were not designated as “Do Not Resuscitate” (DNR), “Comfort Measures Only” (CMO), or “Allow Natural Death” (AND). Our final dataset included 41,707 patients from across 25 hospitals, ranging from large teaching hospitals to smaller, community hospitals [8]).

### Representative samples

When developing a classification algorithm using a supervised training method, one chooses training instances whose labels (i.e., stable versus unstable) are as reliable as possible. However, when evaluating how the algorithm will perform in a real world environment, the test data set needs to be representative of all the types of episodes that it will encounter - ambiguous cases, cases with missing data, very short stays before going unstable, etc. The heterogeneity of such an environment is illustrated by our dataset. It is from a non-annotated database, and as such, no gold standard marker of hemodynamic instability was available. Instead, certain interventions by clinicians were used to demarcate episodes of hemodynamic instability. We developed two lists of intervention criteria for hemodynamic instability - a set of *strong* criteria and a set of *weak* criteria. The first list consisted of intervention criteria indicative of hemodynamic instability for which there was a strong consensus among a group of experienced intensive care physicians (see Table 2). The criteria are differentiated by how specific the intervention is to hemodynamic instability. For example, the administration of norepinephrine is usually a response to hemodynamic instability, whereas the administration of 700cc of fluids in one hour but also could be in response not only to hemodynamic instability, but also in response to other factors. Only the strong criteria were used to define instability when developing the algorithm. One could argue that when validating the only only the strong criteria should be used for defining instability, since the weak criteria are ambiguous and will include many cases which are not really unstable. However, both were used when validating the algorithm, since we wanted to measure how the algorithm would behave in a realistic clinical setting where the algorithm is exposed to all patients throughout their stay, and every alert must be labeled as a true positive or a false positive, even if the actual patient state is somewhat ambiguous.

**Table 2:** Criteria used to define hemodynamic instability were divided into two groups: strong and weak. The strong intervention criteria were used to define the unstable patient population for training purposes, while both the strong and the weak intervention criteria were used for validation of the algorithm’s performance.

| Strong Intervention Criteria | Weak Intervention Criteria |
| --- | --- |
| Administration of any quantity of any of the following inotropic and vasopressor medications:<br>• Dobutamine<br>• Dopamine<br>• Epinephrine<br>• Norepinephrine<br>• Phenylephrine<br>• Vasopressin<br>Administration of Packed Red Blood Cells (PRBCs) in either of the following dosages:<br>• 800 cc PRBC over course of 24 hours<br>• 500 cc PRBC in two hours followed by fluid therapy within 12 hours. (What qualifies as "fluid therapy" is described in the first item the second column of this table, titled "Administration of Fluid Therapy.") | Administration of Fluid Therapy (colloid or crystalloid) in the following dosages:<br>• 700 cc in one hour<br>• 1500 cc total in four hours<br>• 2400 cc in eight hours<br>• 3000 cc in 12 hours<br>• 500 cc twice in four hours<br>Administration of PRBCs in the following dosage:<br>• 500 cc PRBC not followed by fluid therapy within the following 24 hours. (What qualifies as "fluid therapy" is described in the above table entry titled "Administration of Fluid Therapy.")<br>Administration of any of the following pharmacological agents at any dosage:<br>• Nitroprusside<br>• Lidocaine<br>• Amiodarone<br>• Milrinone |

The intervention criteria, both strong and weak, were used to partition each patient’s ICU stay into four types of segments: intervention, pre-unstable, unstable, and stable segments (see Figure 2). The intervention segments were used to define the other segments. An intervention segment began with one of the interventions, weak or strong, listed in Table 2. It continued until there had been no more interventions for 12 hours. In other words, an intervention segment may have included several interventions sustained over various periods of time, followed by 12 hours in which there was no intervention. (Twelve hours was chosen, somewhat arbitrarily, as the time the patient would be under close observation while being “weaned” from the intervention.) An unstable segment was a segment that ended with the initiation of an intervention. Segments that ended with an intervention were divided into unstable and pre-unstable segments. Those that were 24 hours or less in length were labeled unstable segments. Those that were longer than 24 hours were partitioned into the 24-hour period prior to the intervention (an unstable segment), and the period more than 24 hours before the intervention (a pre-unstable segment). Alerts occurring during a pre-unstable segment, i.e. alerts occurring earlier than 24 hours prior to the intervention, were treated as false positives. Alerts occurring during an unstable segment were counted as true positives. Any cutoff time point separating true positives from false positives is going to be somewhat arbitrary. Although the algorithm sometimes detects instability more than 24 hours prior to the intervention, it was decided that alerts earlier than 24 hours were unlikely to be of much help to the clinician. The important point is that in an ICU setting the algorithm is going to face cases like these, and if the goal is to get a realistic evaluation, cases like these cannot simply be ignored. One may decide to handle them differently, e.g., but they need to be handled, and one needs to be explicit about how they are handled.

**Figure 2:**
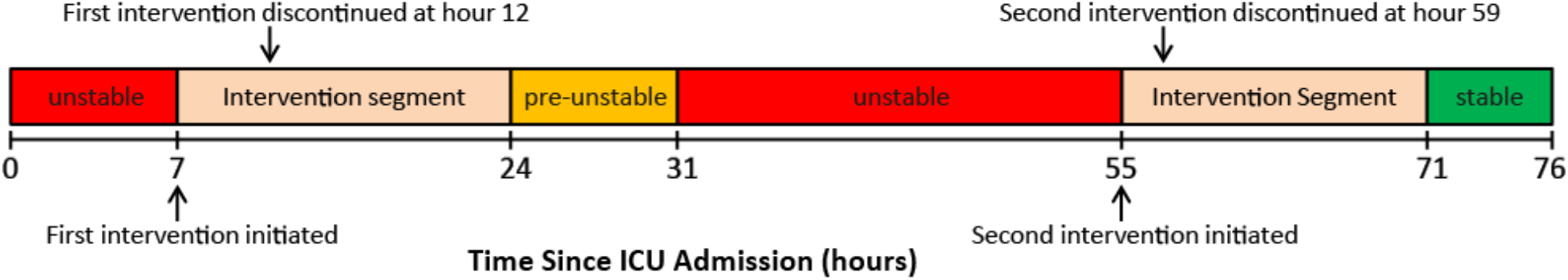
An example of the segmentation scheme used for an ICU patient who stayed in the ICU for a total of 76 hours, and received two interventions. Intervention segments begin during the intervention and end twelve hours after the intervention was discontinued.

Stable segments were all other segments that did not end with an intervention, and any alerts during these segments were treated as false positives. They included the full ICU stay of patients who never had an intervention for the entire encounter and patients who had an intervention but after the intervention segment ended had no additional interventions for the rest of their encounter. For clarity, an example of a segmented patient stay is shown in Figure 2.

In summary, any evaluation if it is to be accurate must include samples that are representative of the actual environment in which it will be running. This includes cases that are not clear-cut either because the labelling is ambiguous (e.g., meet only weak criteria) or are difficult to interpret (e.g., alerts that occur long before the patient becomes obviously unstable). In principle, one could choose sample snapshots that are representative, but a much easier and straightforward solution is to not sample but to use all instances and all data available. But if one is continuously sampling, how does one count instances?

### Consistent counting

Ideally, an algorithm for predicting hemodynamic instability should issue no alerts during the stable and pre-unstable segments and at least one alert during the unstable segments. HII was developed as an indicator whose trend over a specified period of time (e.g., most recent 6 hours) can be displayed to the clinician. So although HII does not issue any alerts, for evaluation purposes it can be treated as an alert - based on its crossing a specified threshold (e.g., greater than 66). Typically, one would evaluate such an algorithm’s performance in terms of its sensitivity, specificity, PPV and NPV, where these are defined in terms of TP, FP, TN and FN. As pointed out in the introduction, these definitions depend upon counting the negative instances (TN and FN) in a way that is comparable to counting the positive instances (TP and FP). But this is not as straightforward as it may seem. Since we are interested in what fraction of the unstable segments that we correctly identify, it is natural to define sensitivity (the true positive rate) as the fraction of unstable segments which have at least one alert. Here a TP in (TP/(TP+FN)) is an unstable *segment* with at least one alert and a FN is an unstable *segment* with no alerts. On the other hand, it is natural to define PPV as the fraction of alerts that are true alerts, since (1-PPV) is essentially a measure of the annoyance factor - how often clinicians are distracted by false alerts. Here a TP in (TP/(TP+FP)) is an *alert* occurring during an unstable segment and a FP is an *alert* occurring during a stable or pre-unstable segment. Thus, in one case we are defining TP in terms of segment counts and in the other case in terms of alert counts.

Specificity and NPV are even more problematic to define. Given how sensitivity is defined, it would seem to follow that the false positive rate (1 - specificity) is the fraction of the combined stable and pre-unstable segments that do not have alerts. But this can be very misleading, as can be seen by reflecting on Table 3: Number and length of segments in evaluation dataset. Note that the stable and pre-unstable segments are considerably longer, on average, than the unstable segments. Since the algorithm is running continuously, this means that there are many more opportunities for a false positive during stable and pre-unstable segments. Intuitively, a five day stable segment with one (false) alert, for example, shouldn’t be given equal weight in the false positive count as a 12 hour stable segment with one (false) alert. The same issue applies to defining NPV.

**Table 3:** Number and length of segments in evaluation dataset

|  | Stable segments | Pre-unstable segments | Unstable segments |
| --- | --- | --- | --- |
| Number of segments | 42,886 | 3,516 | 14,051 |
| Ave length of segments | 2.55 days | 3.22 days | 0.429 days |

So, given all these complications, how do we evaluate an algorithm running in a continuous monitoring environment? We need to do two things. First, we need to divide stable and unstable segments into roughly equal sub-segments. Second, we need to distinguish two ways of parsing positives, one based on counting alerts and the other based on counting sub-segments with alerts. Dividing segments into sub-segments addresses the problem of counting negatives when two segments are of very different lengths. Suppose we choose the sub-segment size to be 24 hours. Then a stable segment that is five days long would have five sub-segments. If a segment is not an exact multiple of 24-hours, then the fraction will also count as a sub-segment. Thus, a stable segment 30 hours long would have two subsegments (6 hours and 24 hours long). Each sub-segment is a potential positive or negative instance, depending upon whether it has at least one alert (positive) or no alerts (negative). The choice of subsegment size is algorithm dependent, and will often depend upon how one wants to treat the detection of the unstable states. If the algorithm is considered successful if it issues a least one alert during the unstable state, then the sub-segment size should be the same length as the maximum length of the unstable segment. On the other hand, if the algorithm is designed to issue multiple alerts during an unstable segment, then the sub-segment size should be chosen so each sub-segment can have one, and only one, alert. For example, the algorithm may include an inhibition period after each alert, so that alerts are suppressed until that period has passed. If the inhibition period is three hours, then setting the sub-segment size to three hours would correspond to counting all possible alerts. Thus, a stable segment four and a half days long would have 36 sub-segments, an unstable segment of 24 hours length would have 8 sub-segments, and one 5 hours long would have 2 sub-segments (2 hours and 3 hours long). This would mean that any of the unstable sub-segments without an alert would count as a false negative, and thus lower the sensitivity, but any of the stable sub-segments without an alert would count as a true negative, and raise specificity.

For the purposes of determining a sub-segment-based sensitivity, specificity, NPV and PPV, the definitions in Table 1 are applied to the sub-segment counts, where each sub-segment is counted as a single positive if there is at least one alert occurring in that sub-segment and as a negative if there is no alert in the sub-segment. Note that this gives us a sub-segment-based PPV that needs to be distinguished from the alert-based PPV.

From the alerting perspective what is analogous to sensitivity is the true positive alert rate (i.e., the number of true positive alerts per unstable patient-day). And what is analogous to 1-specificity is the false positive alert rate (i.e., the number of false positive alerts per stable (and pre-unstable) patient-day).

### PPV and prevalence

Furthermore, just as there are two methods for determining PPV (based on whether one is counting sub-segments or alerts), there are two methods for determining the positive likelihood ratio (pLR). For the alert-based method, the pLR is the ratio of the true positive alert rate to the false positive alert rate. For the sub-segment-based method, the pLR is the ratio of the sub-segment-based sensitivity to the subsegment-based false positive rate (i.e., one minus sub-segment-based specificity). Correspondingly, there are two methods for determining prevalence. The alert-based prevalence is the time duration of all unstable segments divided by the time duration of all segments (stable, pre-unstable, and unstable). The sub-segment method for determining prevalence is the total number of unstable sub-segments divided by the total number of sub-segments. Since, PPV, pLR, and prevalence are closely interrelated, the corresponding alert or sub-segment-based definitions should be used for all three. It is easiest to see this interrelatedness if we convert the prevalence of instability to an odds and invert it. This is the odds of being exposed to stable and pre-unstable episodes versus unstable, which we will refer to as the exposure odds, EO.

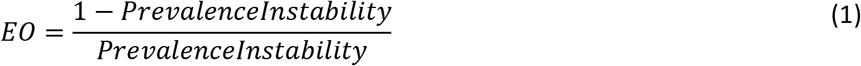

PPV can be calculated as follows:

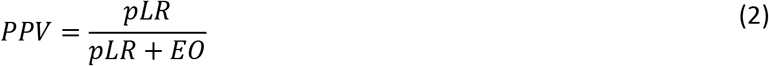

See Appendix for derivation of this relationship.

The results should be the same as when calculated using TP and FP, provided one consistently uses either alert-based or sub-segment-based definitions. This formulation of PPV makes it clear that any measure of PPV is misleading unless it adequately reflects the true prevalence of the state or condition being identified (e.g., instability). This is one of the dangers of using a static “snap shot” analysis. If the static samples are proportional to the stable (control) and unstable (targeted) states, they can be used to get an accurate estimate of sensitivity and specificity. But unless they reflect the true prevalence, i.e., the ratio of number of stable snapshots to unstable snapshots equals the true exposure odds, a PPV directly calculated from the snapshots will not represent the true PPV. It needs to be adjusted to reflect the prevalence. This follows from equation (2).

Although we haven’t given much attention to NPV, the sub-segment version of NPV can easily be calculated directly from the sub-segment counts, or indirectly using the negative likelihood ratio (nLR) and EO:

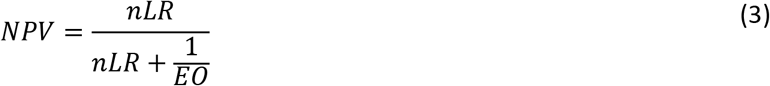

where *nLR* = *specificity*/(*1-sensitivity*).

Furthermore, an alerting version of NPV can also be calculated. In this case one must use the alert-based EO, and calculate nLR by subtracting both the true positive alert rate and false positive alert rate from the maximum possible alerts per day, and taking the ratio. For example if there is a three hour inhibition period after an alert, then the maximum number of alerts per day is eight, and nLR = (8 - FP_a lert_rate)/(8 - T P_a lert_rate).

### Dynamic alert supervision

As noted earlier, in the case of the HII algorithm, when determining sensitivity, we were concerned about the fraction of unstable segments that we caught, not whether we caught them multiple times, so we used a sub-segment size that was the same size as the maximum length of unstable segments, 24 hours. Furthermore, to simulate how a clinician might react to an indicator whose trend is being displayed, we used a dynamic alert supervisor. The dynamic alert supervisor disarms further alerts, once an initial alert has been issued, only re-arming alerts when HII had significantly deteriorated. For example, in response to the first time that the indicator goes into the “danger” zone, the clinician reviews the patient’s situation. The clinician determines that the patient is not actually in danger and does not require an intervention, perhaps because this patient has a low baseline blood pressure. The clinician is unlikely to continue giving this patient the same amount of attention as the first time it went into the “danger” zone, unless the indicator significantly worsens. For evaluating HII, significant deterioration was defined as half the sum of 100 and the HII value when the previous alert was issued. For instance, if the first alert was issued when HII had a value of 70, no additional alerts were issued until HII was greater than 85 (i.e., (100+70)/2), and the three hour inhibition period had been met. If a second alert is issued when HII has a value of 88, a third alert will not be issued unless HII has a value greater than 94 (i.e., (100+88)/2). So once an alert is issued, the number of possible subsequent alerts can only be determined dynamically.

There are many other possible ways of dynamically supervising alerts. One might, for example, require after any alert that there is a sustained period without any alert threshold crossing before a subsequent alert is issued. Dynamic alerting schemes make it difficult to evaluate alerting behavior using a snapshot approach, since what happens in a snapshot may depend on what happens in previous snapshots.

## RESULTS

As we noted in the introduction, there are many ways to evaluate an algorithm such as HII. One approach is to take a “snapshot” approach, looking at discrete data points chosen from stable and unstable segments. We are not arguing here that this approach should not be used. It is quite useful when comparing the relative performance of two algorithms on the same set of snapshots. We used this approach to measure how early HII detected hemodynamic instability in the ICU before the onset of acute hypotension. [6] This allows us to use area under the receiver operator curve (ROC) analysis to measure sensitivity-specificity tradeoffs, comparing HII’s performance with an algorithm that simply uses systolic blood pressure as the basis for alerts. Figure 3 shows how the area under the ROC curve (AUC) changes for a range of prediction windows, comparing HII with SBP, treating HII and SBP as continuous variables rather than alerts. As one can see from the plots HII outperforms SBP. Furthermore, for a prediction window of 12 hours, HII still has considerable predictive value, whereas SBP has little predictive value (an AUC of 0.5 would indicate zero predictive value).

**Figure 3:**
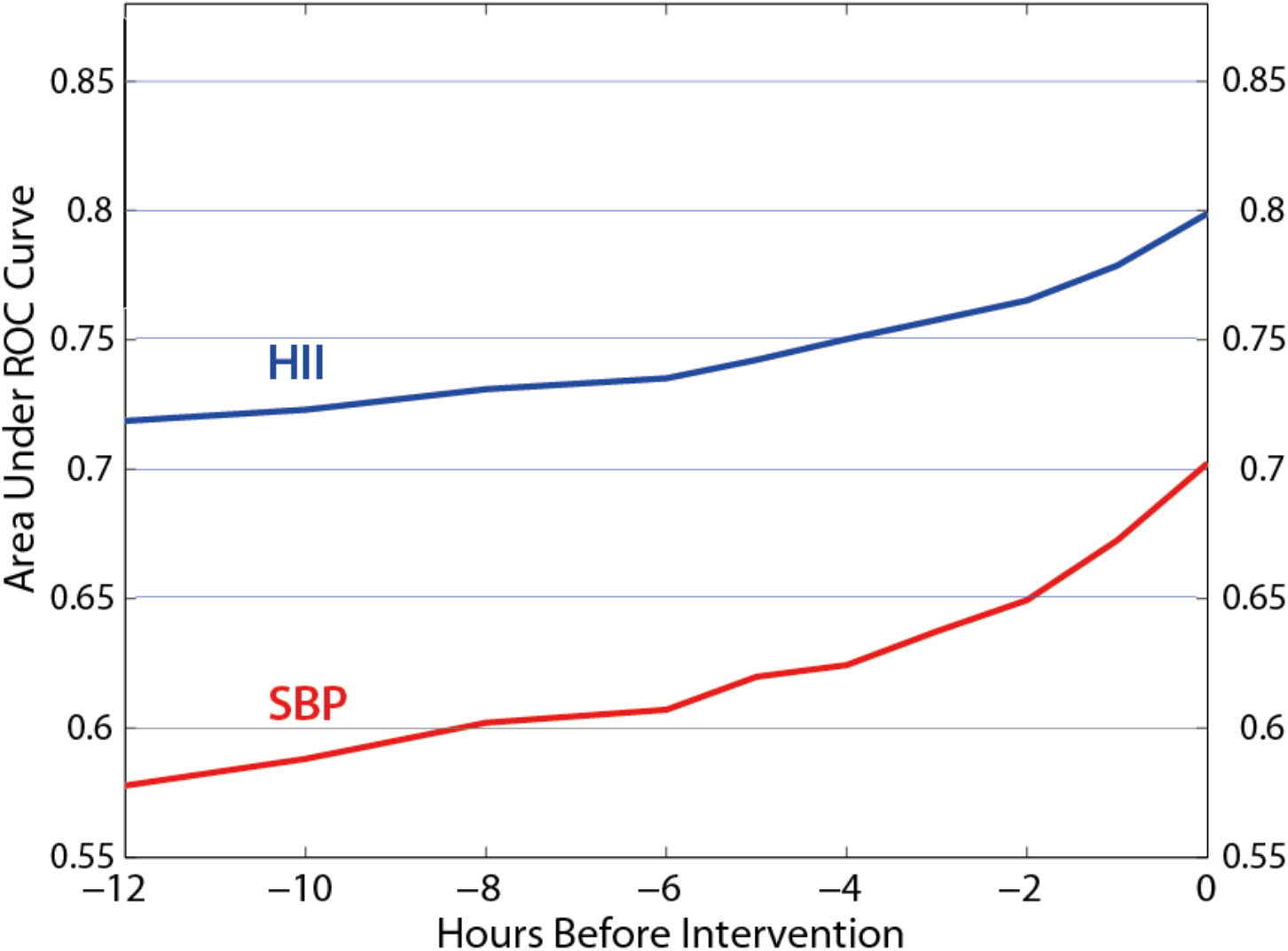
AUC for prediction windows ranging from 0 hours to 12 hours prior to intervention for measured SBP and the corresponding HII value.

But one cannot infer from such an analysis how HII will behave in a real ICU setting since the snapshots in this analysis are not representative. It only uses data from unstable segments that are greater than 12 hours long and HII and SBP are available throughout the segment, but only 26% of unstable segments fit these criteria.

This diversity in unstable segment size (Figure 4) is going to affect performance, but it is difficult to appreciate how by simply analyzing snap shots. For example, an unstable segment that is 24 hours long is much more likely to trigger an alert than an unstable segment that is one hour long. A snapshot-based analysis will give us some interesting insights, but there are many questions that can only be answered by simulating performance when running continuously. How effective will it be at identifying unstable episodes when these episodes are of varying length? What will the alert burden on the clinician be?

**Figure 4:**
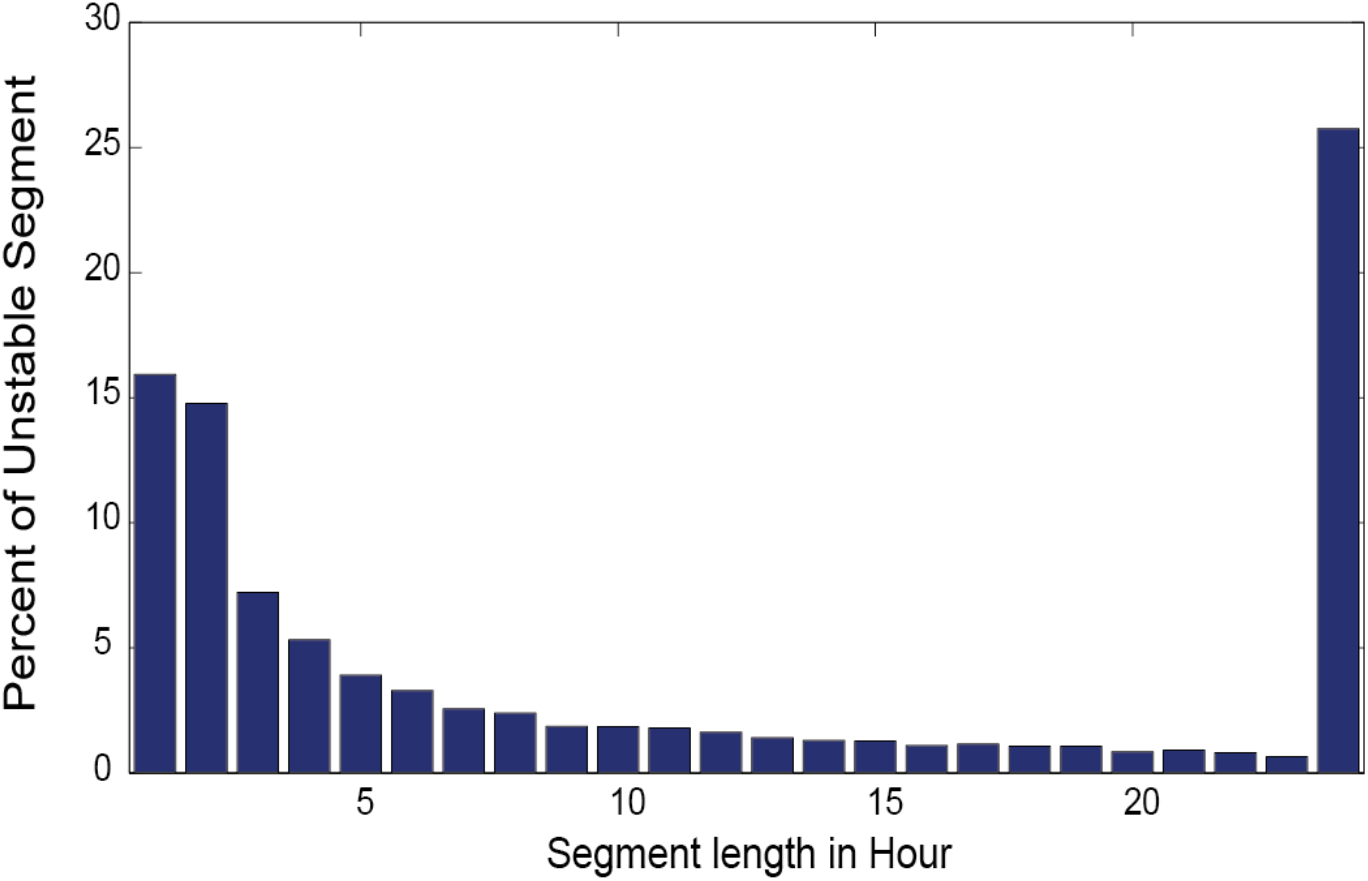
Distribution of unstable segment lengths. The maximum length of unstable segments is capped at 24 hours, hence the peak at 24 hours.

To answer these sorts of questions we simulated HII’s performance in an ICU setting on the retrospective dataset, consisting of 41,707 patients from across 25 hospitals’ ICUs from the eRI dataset. All the unstable, pre-unstable, and stable segments were used for final validation purposes. It is important to emphasize that these results represent performance on real-world ICU data; there was no pre-screening of the data. All patients in the dataset were included in the analysis. The dataset consisted of 14,051 unstable (potentially TP) segments and 46,402 stable and pre-unstable (potentially FP) segments (Table 3). Heart rate and arterial line systolic blood pressure (SBP) were reported as the median values for consecutive, five-minute non-overlapping intervals. When arterial blood pressure was not available, noninvasively measured systolic blood pressure was used for both HII and the SBP alerts.

For alerting purposes a HII threshold of 66 (>) was used which puts HII is the top third of its range. For SBP alerts several threshold were tried. Here we show results for a threshold of 80 mmHg (<), which is a fairly common (if somewhat conservative) alerting threshold. The “unsupervised” versions of both algorithms used a 3-hour inhibitory period after each alert. The “supervised” versions, in addition to the 3-hour inhibitory period, suppressed all subsequent alerts, after the initial alert, unless there was a significant deterioration. In the case of HII this was defined as half of the sum of 100 and the HII value when the previous alert was issued. In the case of SBP a significant deterioration was defined as half the sum of 40 mmHg and the SBP value when the previous alert value was issued.

The exposure odds in terms of alert exposure, i.e., the ratio of time exposed to false positive segments (stable and pre-stable segments) to time exposed to true positive segments (unstable segments), was 20.0 (i.e., a prevalence of 0.0476). The exposure odds in terms of 24 hour sub-segment counts, i.e., the number of stable and pre-unstable sub-segments divided by the number of unstable sub-segments, was 10.15. Table 4 shows the results for HII and SBP using a 24-hour sub-segment period (the same size as the maximum unstable period). The ssSensitivity (sub-segment-based Sensitivity) provides a good estimate of what fraction of unstable episodes will be caught by various algorithms in a realistic simulation of an ICU setting, and the aPPV (alert-based PPV), as well as the FP alert rates, provides a good estimate of the annoyance factor due to false alerts. Table 4 indicates that the supervised version of HII has a considerable advantage over the supervised version of SBP (dominating on both sensitivity and aPPV) and the unsupervised version of HII has a slight advantage over the unsupervised version of SBP (dominating on sensitivity when aPPV are about the same).

**Table 4:** Comparison of HII to SBP as an alerting algorithms with respective thresholds of HII > 66 and SBP < 80 mmHg for versions with (Sup) and without (Uns) dynamic alert supervision. The prefix ‘ss’ indicates measurements based on sub-segments (size 24 hours). aPPV is the alert-based PPV. TP and FP alert rates are reported for alerts per patient day.

| Algorithm | ssSpecificity | ssSensitivity | ssPPV | FP alts/pday | TP alts/pday | aPPV |
| --- | --- | --- | --- | --- | --- | --- |
| HII>66,<br>Unsupervised | 0.882 | 0.388 | 0.245 | 0.332 | 1.859 | 0.219 |
| HII>66,<br>Supervised | 0.934 | 0.332 | 0.338 | 0.079 | 0.821 | 0.341 |
| SBP<80,<br>Unsupervised | 0.881 | 0.349 | 0.224 | 0.197 | 1.134 | 0.223 |
| SBP<80,<br>Supervised | 0.919 | 0.303 | 0.269 | 0.097 | 0.714 | 0.269 |

Interestingly, the ssPPV (sub-segment-based PPV) is quite close to the aPPV (alert-based PPV) in three of the algorithms. This will happen when the number of alerts per unstable sub-segments with alerts is almost the same as the number of alerts per stable and pre-unstable sub-segments with alerts. This is the case when dynamic alert supervision is used. Most sub-segments that have an alert do not have a subsequent ones because the dynamic alert supervision mechanism inhibits all subsequent alerts that do not reflect a worsening condition. Thus alert supervision reduces alert rates of the unstable segments to a level that is close to the low alert rate of the stable and pre-unstable episodes. Therefore, it is not surprising that the aPPV and ssPPV are nearly the same for both algorithms that use dynamic alert supervision. In the case of the algorithms without dynamic supervision, sub-segments often have multiple alerts. Given the higher alert rate for unstable segments than other segments, one might expect that the average number of alerts per unstable sub-segment with alerts would be higher than the FP (stable and pre-unstable) sub-segments. If this were the case, then the aPPV would be higher than the ssPPV. But as Table 4 shows, this is not the case. In fact, for HII the ssPPV is higher than the aPPV. Two things are happening that explain this. First, although the unstable segments have a higher alert rate than the FP segments, they are on average much shorter (see Table 3: Number and length of segments in evaluation dataset). In fact 38% of the unstable segments are three hours or less in length and so cannot have more than one alert due to the three hour inhibition period. It just so happens that in the case of the unsupervised SBP algorithm the average number of alerts per FP sub-segments with alerts is almost the same as for the average number of alerts for unstable sub-segments with alerts, whereas in the case of the unsupervised HII algorithm the average is higher in FP sub-segments than the unstable sub-segments. And this brings us to the second thing that is happening. The HII algorithm predicts instability much earlier than the SBP algorithm. This early alerting often extends beyond the unstable segments to the pre-unstable sub-segments, which unlike the unstable sub-segments, often are of maximum length, and so often have multiple alerts. Because HII has a higher number of FP pre-unstable alerts than the SBP algorithm, the average number of alerts per FP sub-segment is higher than the average number of alerts per unstable sub-segment, and thus the aPPV is lower than the ssPPV. This is a good example of where the dynamics of an algorithm can be quite subtle and can be missed when one simply looks at static snapshots. Although HII predicts hemodynamic instability much earlier than the SBP algorithm, this advantage is partially offset by the predominance of short unstable segments and a higher FP rate for the pre-unstable segments.

The results in Table 4 were based on sub-segments the same size as the maximum unstable segment (24 hours). Table 5 shows the results when a 3-hour sub-segment period is used. (Since the alert rates are not affected by the sub-segment size. Comparing Table 4 and Table 5, one will note that the ssPPV is now the same as the aPPV. This is because the alerts have a 3-hour inhibition period, which is the same as the sub-segment size, so there can only be one alert per sub-segment. The big difference, however, is that sensitivity is much lower for the algorithms shown in Table 5. This is because unstable segments longer than 3 hours are treated as multiple sub-segments, with up to 8 sub-segments in a stable segment. If any of these sub-segments does not have an alert, they are counted as false negatives in the sensitivity calculation. Furthermore, the drop in sensitivity is especially large for the supervised versus unsupervised version of HII. This reflects the fact that HII tends to detect instability quite early, and if the unstable segment is longer than 3 hours, it often will detect instability in one of the earlier subsegment (i.e., earlier than 3 hours before the intervention), but the alert supervisor will suppress subsequent alerts if there is not significant deterioration and so the later unstable sub-segments will count as a false negatives. Finally, the sensitivity is much lower for the unsupervised version of SBP than the unsupervised version of HII. Again, this reflects the fact that HII detects instability earlier and thus earlier 3-hour unstable sub-segments are more likely to have alerts in the case of HII than the SBP algorithm. (For 3-hour sub-segments the exposure odds are 18.4.)

**Table 5:** Same as Table 4, except the sub-segment size used is 3 hours instead of 24 hours.

| Algorithm | ssSpecificity | ssSensitivity | ssPPV | FP alts/pday | TP alts/pday | aPPV |
| --- | --- | --- | --- | --- | --- | --- |
| HII>66,<br>Unsupervised | 0.959 | 0.209 | 0.219 | 0.332 | 1.859 | 0.219 |
| HII>66,<br>Supervised | 0.990 | 0.092 | 0.341 | 0.079 | 0.821 | 0.341 |
| SBP<80,<br>Unsupervised | 0.976 | 0.127 | 0.223 | 0.197 | 1.134 | 0.223 |
| SBP<80,<br>Supervised | 0.988 | 0.080 | 0.269 | 0.097 | 0.714 | 0.269 |

In summary, we applied a new method of evaluating performance to the HII algorithm and compared to SBP alerts in predicting hemodynamic instability. The proposed methodology allowed us to characterize these algorithms’ behavior in a more complete way than traditional method allows. From the snapshot approach alone, we know that HII outperforms SBP in detecting instability earlier, but the analysis only used 26% of unstable episodes. While useful, this does not give us a sense of how the algorithm would perform on a more heterogeneous dataset (as found in real ICUs), nor the alert fatigue that the algorithm would impose on clinicians. Using the sub-segment approach and taking carefully defining positive and negative instances, we are able to capture more subtle but important characteristics of the algorithms: 1) We calculate the sensitivity in detecting unstable patient episodes regardless of how brief the patient has been in the ICU, 2) we quantified the reduction in alert fatigue that dynamic supervision would bring, and 3) the ability of HII to predict instability very early may have negative implications if clinicians find this too early to be useful.

## DISCUSSION

The sub-segment approach can be looked upon as a dynamic version of the snapshot approach. One is taking a continuous series of snapshots (as in a moving picture) throughout the patient’s stay. There are several advantages of the dynamic approach. By sampling continuously and throughout each patient’s stay one does not need to worry about the samples being representative or reflecting the actual population prevalence. Furthermore, it forces one to make a decision about how large the sampling window (i.e., sub-segment size) should be, depending upon what one is trying to measure. A maximum sub-segment size equal to the maximum instability segment size, for example, will measure how many unstable segments have at least one alert. Also, one is not limited to choosing the sub-segment size to be either equal to the maximum unstable segment size or the alerting inhibition period. Depending on one’s purposes one may choose some alternative such as half the maximum sub-segment size, or explore several alternatives. The alert-based approach, on the other hand, nicely measurers the annoyance factor of too many false alerts. There is no requirement that one choose between the two approaches to counting. One can use both approaches. What is important is that one recognizes that they are different approaches with different methods of counting positives and negatives.

The methodology introduced provides a flexible way of analyzing performance of an indicator or alerting algorithm that is being run continuously with the potential for many positives (true or false) on the same patient.

## CONCLUSION

Given that ICU clinicians are already suffering from alert fatigue, when we created HII our first priority was to design an algorithm that does not significantly add to this burden. To accurately assess HII’s performance, including how it might add to the clinician’s alert burden, we developed a methodology for evaluating HII’s performance using retrospective data that simulates how HII might perform in a real ICU setting. We wanted the algorithm to be exposed to situations that were representative, reflected the population prevalence of hemodynamic instability episodes, and reflected realistic alerting dynamics. The algorithm was assumed to be running continuously throughout the patient’s stay (except during interventions). All patients were included except those designated “Do Not Resuscitate”, “Comfort Measures Only”, or “Allow Natural Death”. Some of the unstable segments included in this study occurred in patients who were administered an intervention for hemodynamic instability documented shortly after they entered the ICU; in these cases, an insufficient window of time was present for the algorithm to generate an alert. Such cases were, nonetheless, included in the retrospective validation of the algorithm, as they represented real clinical scenarios. Furthermore, to get an accurate assessment of the alerting behavior of the algorithm, both strong and weak criteria were used for classifying stable and unstable segments. These are based on commonly utilized interventions in the ICU, but are certainly not a “gold standard” for labeling a patient as truly stable or unstable. In fact, the weak criteria indicate situations where clinicians often disagree as to whether interventions are warranted. Likewise, the 24-hour cut-off for instability and the inclusion of earlier alerts as false positives is an arbitrary dividing line, but was made necessary by our desire to make our evaluation of performance as realistic as possible. Although we believe that our methodology for evaluating algorithms like HII on retrospective data provides a realistic basis for predicting how HII will *behave* in a prospective clinical study, only an actual prospective study can determine whether clinicians find this behavior of value. Based on its behavior on retrospective data, we can predict, for example, that HII will detect instability in many patients very early (sometimes more than 24 hours early), but only an actual clinical study can determine if clinicians find this behavior useful or annoying. Previous studies have certainly shown that earlier detection and timely intervention can significantly improve patient outcome [3, 4, 5].

Continuous monitoring is becoming more and more common, not only in the ICU, but in the emergency department, the general ward, and even in the home (telemonitoring). Alerting and detection algorithms, including acuity indicators, developed for the continuous monitoring environment can only be properly evaluated by recognizing how this environment differs from the more familiar diagnostic testing environment. The proposed methodology makes explicit the underlying assumptions and provides a flexible and customizable approach to evaluating algorithm performance when running continuously.

## COMPETING INTERESTS, FUNDING

Philips is an international, publicly traded, for-profit company. This research was self-funded.

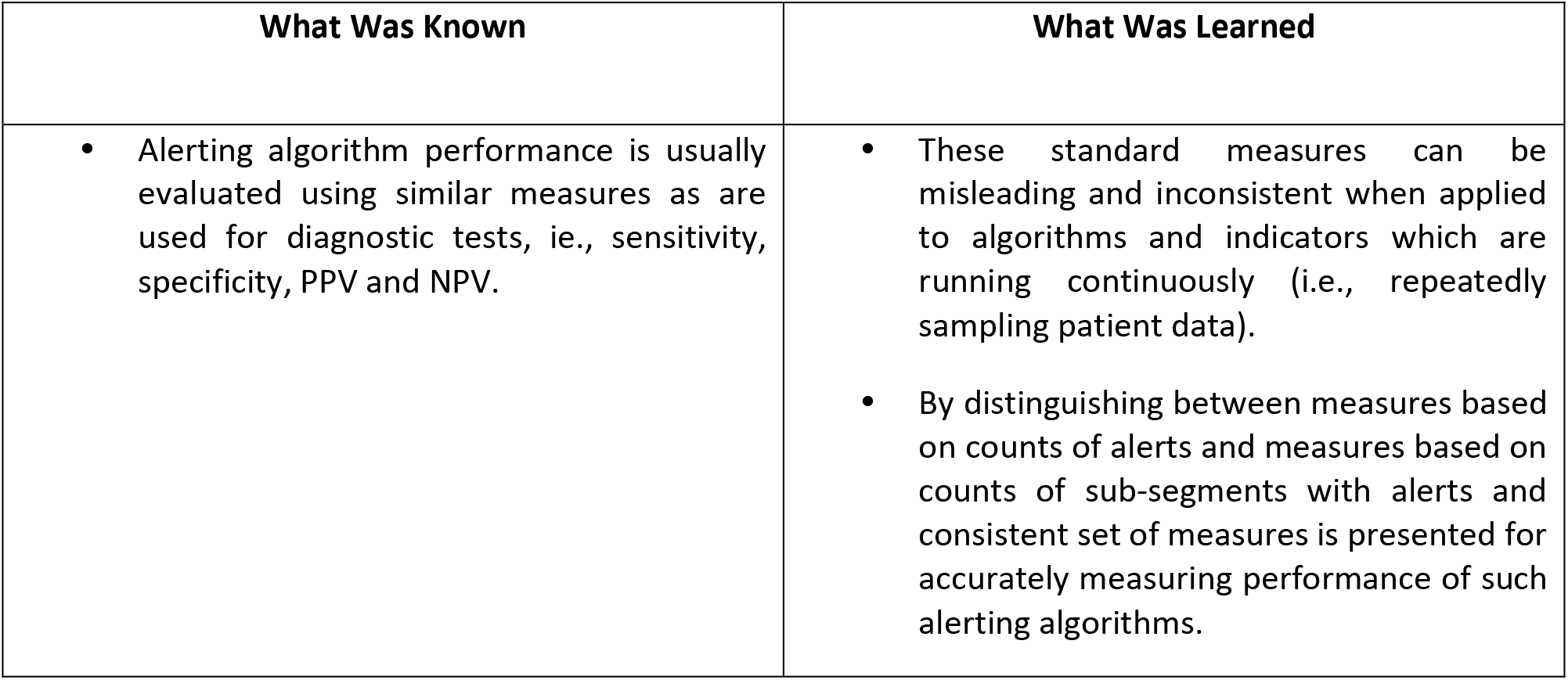
SUMMARY TABLE

## Appendix: Relationship between PPV, pLR, and prevalence

Let

~~~
        Pr(*u*) = *probability of a episode being unstable*
        Pr(*s*) = *probability of a episode being stable*
   Pr(*A*|*u*) = *probability of an unstable episode having an alert*
   Pr(*A*|*s*) = *probability of a stable episode having an alert*
~~~

Then PPV is simply Pr(*u*|*A*). By Bayes’ Theorem,

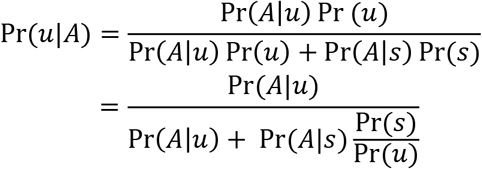

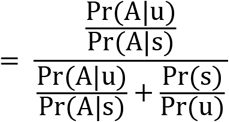

Therefore

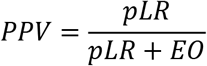

where 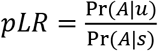 is the likelihood ratio and 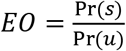 is the exposure odds, the fraction of time when the algorithm is running on stable episodes versus unstable episodes. This is the reciprocal of what is normally called prevalence. As is well-known, PPV is not an intrinsic property of a test, but rather depends on the prevalence of the population. If only a small fraction of an ICU population ever becomes unstable (or only for a small fraction of the time), then *EO* » 1 and one cannot hope to achieve a high PPV.

